# Recombinant expression of Proteorhodopsin and biofilm regulators in *Escherichia coli* for nanoparticle binding and removal in a wastewater treatment model

**DOI:** 10.1101/286195

**Authors:** Justin Yang, Yvonne Wei, Catherine Yeh, Florence Liou, William Chen, Dylan Lu, Christine Chen, Justin Pei, Candice Lee, Emily Chen, Ashley Lin, Paul Imbrogulio, Katie Chang, Andrew Hu, Jesse Kao, Kelly Chen, Audrey Tei, Chansie Yang, Katherine Hsu, Laurent Hsia, Oscar Wallace, Abby Hau, Allen Liu, William Huang, Stephanie Chang, Catherine Chang, Leona Tsai, Avery Wang, Chang Sun Lee, Alvin Wang, Moksha Shah, Leon Yim, Sean Tsao, Teresa Chiang, Jude C. Clapper

**Affiliations:** Taipei American School, Taipei City, Taiwan

## Abstract

The small size of nanoparticles is both an advantage and a problem. Their high surface-area-to-volume ratio enables novel medical, industrial, and commercial applications. However, their small size also allows them to evade conventional filtration during water treatment, posing health risks to humans, plants, and aquatic life. This project aims to remove nanoparticles during wastewater treatment using genetically modified *Escherichia coli* in two ways: 1) binding citrate-capped nanoparticles with the membrane protein Proteorhodopsin, and 2) trapping nanoparticles using *Escherichia coli* biofilm produced by overexpressing two regulators: OmpR234 and CsgD. We demonstrate experimentally that *Escherichia coli* expressing Proteorhodopsin binds to 60 nm citrate-capped silver nanoparticles. We also successfully upregulate biofilm production and show that *Escherichia coli* biofilms are able to trap 30 nm gold particles. Finally, both Proteorhodopsin and biofilm approaches are able to bind and remove nanoparticles in simulated wastewater treatment tanks. We envision integrating our trapping system in both rural and urban wastewater treatment plants to efficiently capture all nanoparticles before treated water is released into the environment.

**Financial Disclosure:** This work was funded by the Taipei American School. The funders had no role in study design, data collection and analysis, decision to publish, or preparation of the manuscript.

**Competing Interests:** The authors have declared that no competing interests exist.

**Ethics Statement:** N/A

**Data Availability:** Yes – all data are fully available without restriction. Sequences for the plasmids used in this study are available through the Registry of Standard Biological Parts. Links to raw data are included in Supplementary Information.

## Introduction

### 1.1 Nanoparticle Applications and Potential Risks

Nanoparticles (NPs) are generally defined as matter measuring 1 to 100 nm in at least one dimension [1]. The small size and high surface-area-to-volume ratio of NPs make them ideal for novel applications in many fields such as medical imaging, drug delivery, cosmetics, clothing, personal care and filtration. Currently, the Woodrow Wilson International Center of Scholars and the Project on Emerging Nanotechnologies (PEN) lists around 2000 consumer products containing nanomaterials in over 20 countries, with silver, carbon, titanium, silicon, zinc and gold being the most common materials used in products [2].

The recent increase of nanomaterial usage in consumer products has led to higher exposure rates. NPs are most commonly used in health and fitness products [2], including sportswear, sunscreens and cosmetics. For instance, Silver nanoparticles (AgNP) is a common coating on antimicrobial sports fabrics [3]; however, AgNPs have been shown to be released from fabrics when incubated in artificial sweat [4]. This suggests that AgNPs can fall out during exercise, making dermal exposure to AgNPs very likely. TiO_2_ and ZnO NPs are often used as the primary UV blocking agent in sunscreens (figure 1) because of their transparent appearance, smooth application and broad spectrum UV protection [5]. These products are applied topically and worn for long periods of time, increasing the likelihood of NP exposure.

**Figure 1.**
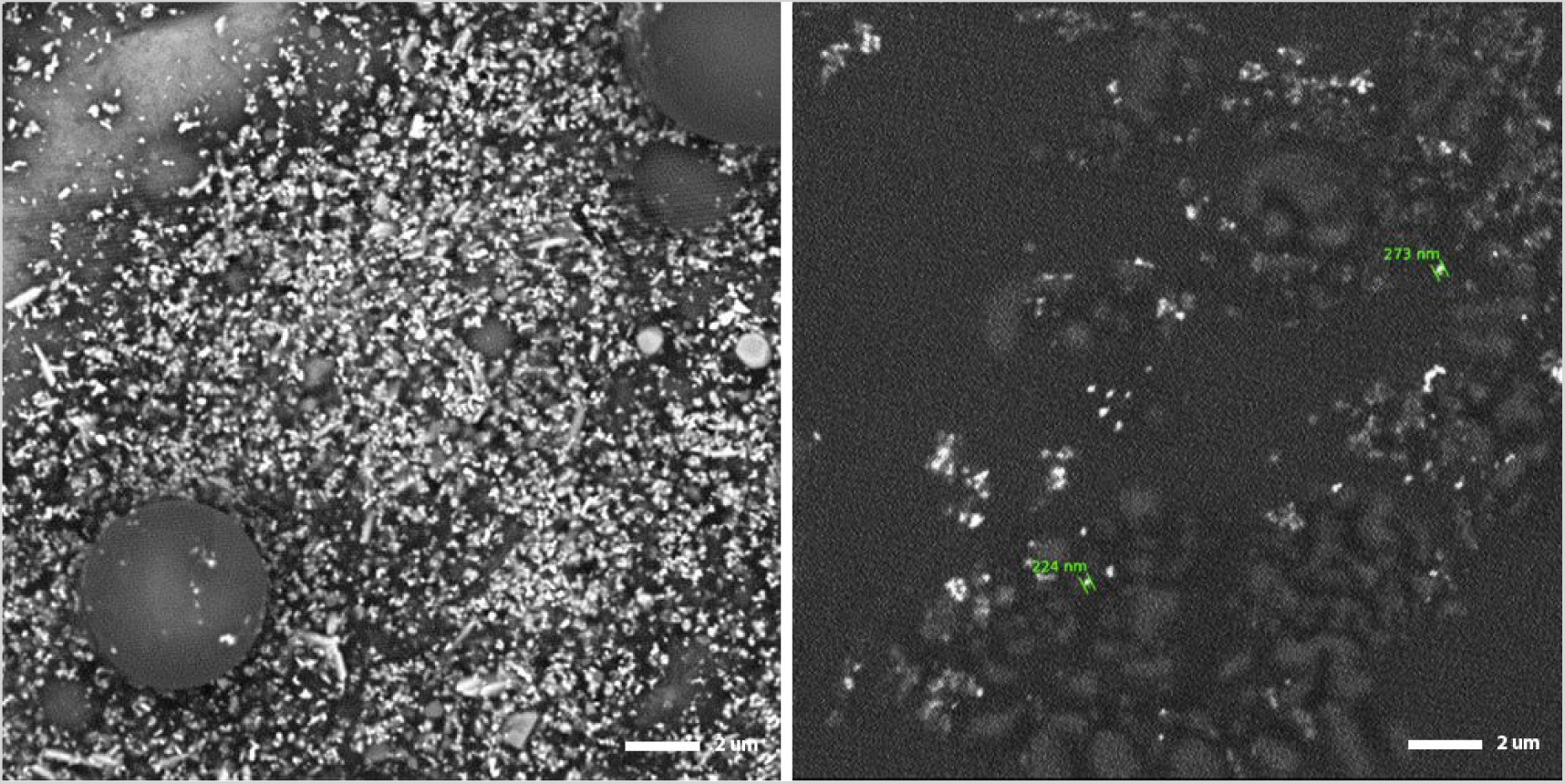
SEM imaging of foundation (cosmetics; left) and sunscreen (right) reveals bright white dots, which indicates the presence of NPs.

The small size of NPs makes them useful in consumer products, but also more reactive and often more toxic than larger, bulk-sized chemicals. When tested in *E. coli*, toxicity dramatically increased when exposed to small silver particles under 10 nm [6]. In another study with crustaceans, algae and protozoa, CuO NPs were shown to be up to 5000 times more toxic than microparticles (i.e. particles between 0.1 µm and 100 µm) [1, 7].

While long term effects of NPs exposure are largely unknown due to the relative novelty of nanotechnology, numerous *in vitro* and *in vivo* studies point to potentially negative health and environmental effects. ZnO NPs can inhibit root growth in common wetland plant species [8]. Fathead minnow (*Pimephales promelas*) embryos experience death or growth abnormalities after exposure to various concentrations of AgNPs [9]. Furthermore, *in vitro* studies have reported negative effects on human cells. Paddle-Ledinek *et al.* found that antimicrobial wound dressings containing AgNPs are cytotoxic to skin cells (keratinocytes); the authors noted “disordered” morphology and decreased cell proliferation, viability and metabolism just 3 hours after exposure [10]. 20 nm AgNPs have also been shown to have detrimental effects to neuronal development; exposure reduced cell viability of premature rat neurons and triggered degeneration of mature rat neurons [11].

### 1.2 Current Nanoparticle Removal in Wastewater Treatment Plants

As nanotechnology becomes an integral part of our daily lives, nanomaterials and their wastes are expected to enter—and likely are already polluting—our natural environment due to inadequate disposal methods. It is estimated that about 95% of AgNPs and TiO_2_ NPs used in consumer products end up in wastewater [12]. Considering that most households in the US are connected to public sewers, Holder *et al.* estimates that up to 2.7 tons of AgNPs and 229.3 tons of TiO_2_ enter municipal wastewater treatment plants (WWTPs) annually in the US [13].

Despite the need to prevent NP pollution, current municipal WWTPs do not have specialized procedures to remove NPs in wastewater. Instead, NPs are subject to the conventional procedures used to treat larger particulates (figure 2). When wastewater enters a plant, a grit screen first removes coarse solids and large materials. The water then moves into a primary sedimentation tank, where heavy solids are removed by sedimentation while floating materials (such as oils) can be taken out by skimming. However, dissolved materials and colloids—small, evenly dispersed solids such as NPs—are not removed here [14]. Next, aerobic microbes help to break down organic materials in aeration tanks; this is also known as the activated sludge process [15]. In a subsequent flocculation and sedimentation step, the microbes are removed and the effluent is disinfected (often by chlorine or UV) before it is released into the environment.

**Figure 2.**
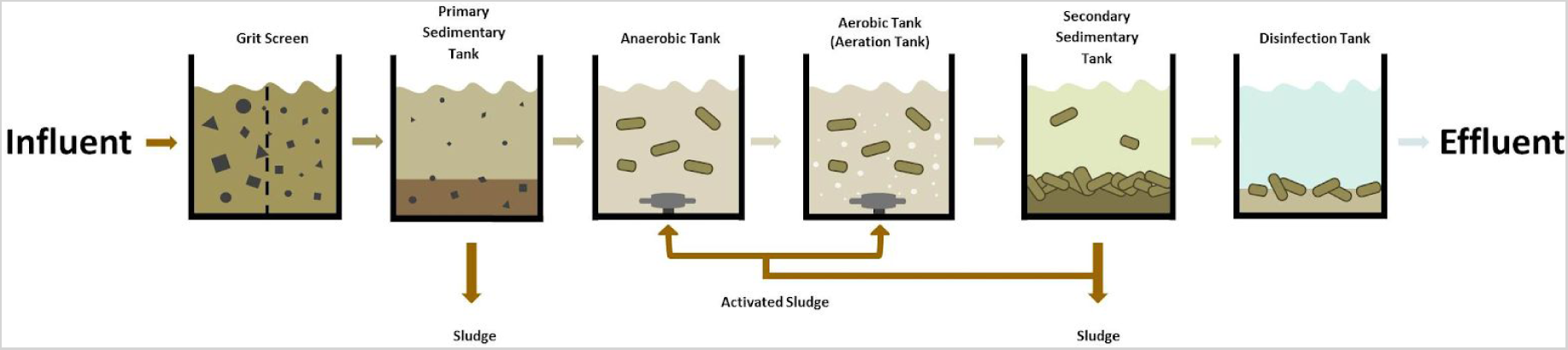
Typical wastewater treatment process. Large particles are removed by filtration and sedimentation, and organic materials are broken down by microbes added to anaerobic and aerobic tanks. Then, a secondary sedimentation stage precedes disinfection before the effluent is released.

While these processes can remove some NPs, complete NP removal has not yet been achieved. A study monitoring an Arizona WWTP found that while sedimentation is effective at filtering large aggregates (72% removal rate), most small-sized TiO_2_ NPs (41% removal rate) can still pass through the WWTP and enter major water systems downstream [16]. SEM images of treated effluent from our local WWTP also contain NPs (figure 3).

**Figure 3.**
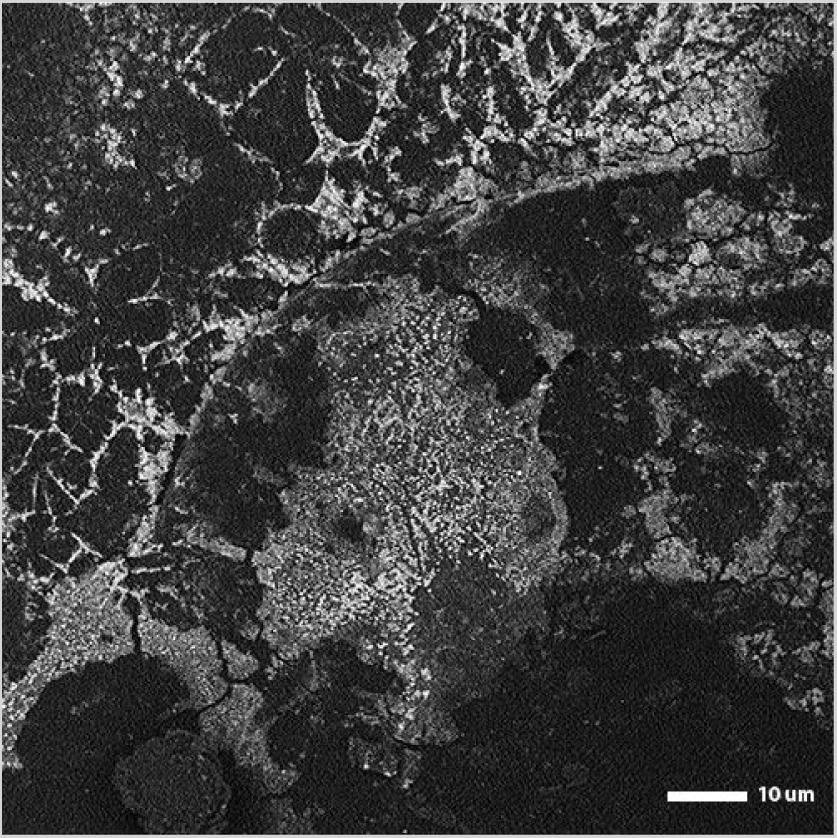
Effluent water from Dihua WWTP contains NPs. Effluent samples from our local WWTP, Dihua, were imaged using SEM. We observed NPs (bright white dots) in these sample, which means that NPs were not completely removed in the wastewater treatment process.

### 1.3 Proposed Nanoparticle Removal Strategies

Our goal is to efficiently remove NPs from wastewater systems to prevent NP pollution. The varying types and sizes of NPs, however, make this task difficult. Approaches that exploit highly specific properties of one type of NPs are inefficient, because they will not work for other NPs. Here, we describe a two-pronged approach to maximize the capture of all NP types.

#### Proteorhodopsin

Most NPs found in consumer products or created for research have a “coating”—known as capping agents—on their surface to prevent aggregation. This makes capping agents a unifying property across different types of NPs. Citrate is the most common capping agent used by industry [17], and a membrane protein called Proteorhodopsin (PR), found in marine proteobacteria, is capable of binding to citrate [18]. **Our goal is to express PR in *E. coli* to bind and hold onto citrate-capped NPs (CC-NPs).**

Ideally, we would add our PR bacteria into existing aeration tanks at WWTPs, where the steady influx and mixing of air provide oxygen favorable to aerobic microbes; the turbulent water also increases the probability of PR binding to CC-NPs. Like many other microbes used in aeration tanks, PR bacteria should increase in size and weight after it captures NPs, such that existing infrastructure in WWTPs can filter them out before NPs get released into natural water bodies.

#### Biofilm

Most WWTPs already use microbes to break down and remove organic components in wastewater [19]. The recent increase in NP contaminants entering WWTPs, however, poses a threat to these microbes and the treatment process. For example, AgNPs have antimicrobial effects and other metal oxide NPs can inhibit microbes from performing important processes such as nitrification [20, 21]. Biofilms are communities of microbes embedded in a matrix of extracellular polymeric substances (EPS), which consist of different polysaccharides, proteins, and lipids [22]. Recent studies show that the EPS in biofilms can trap various NPs [23] and are nearly four times more resistant to NPs, increasing their tolerance to NPs in wastewater [24].

Our preliminary tests and literature research show that biofilms trap NPs and pull them out of solution. *E. coli* produces biofilms through the curli operon, which is regulated by two proteins, OmpR and CsgD. There are many other regulatory mechanisms regulating *E. coli* biofilm synthesis, but since biofilm formation is commonly associated with diseases such as urinary tract infections (UTIs), we avoided genes associated with virulence [25]. Using a safe and common laboratory strain— *E. coli* K-12—as our chassis [26], **our goal is to increase biofilm yield by constitutively overexpressing OmpR and CsgD**. Biofilms can then be used in secondary sedimentation tanks, where the water is relatively calm to keep biofilm structures intact. In addition, larger particles in wastewater would already be filtered out by this stage, which can maximize NP capture rate.

## Materials and Methods

### 2.1 Proteorhodopsin

#### Construct Design

The open reading frame (ORF) of *pR* [18] was modified to remove three internal cutting sites (EcoRI, PstI, and SpeI). The sequence was flanked by a strong promoter and strong ribosome binding site (RBS) combination (BBa_K880005), and a double terminator (BBa_B0015) to maximize PR protein expression. This final construct (BBa_K2229400; figure 4) was ordered from Integrated DNA Technologies, cloned into pSB1C3 (a biobrick backbone), and sequenced by Tri-I Biotech.

**Figure 4.**
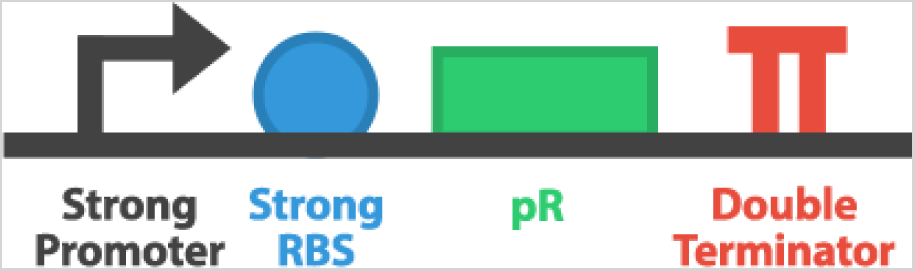
Construct design for PR expression (BBa_K2229400) includes a strong promoter, strong RBS, the *pR* ORF, and a double terminator.

#### Proteorhodopsin and Citrate-Capped Silver Nanoparticles Binding Assay

We tested PR’s ability to trap 60 nm CC-AgNPs (Sigma Aldrich, 730815). Two groups were set up: *E. coli* carrying BBa_K2229400 (PR expression construct) or a negative control BBa_E0240 (GFP-generator construct) were grown in LB overnight. Cultures were centrifuged and resuspended in distilled water to standardize the bacterial populations, then mixed with 0.001 mg/mL CC-AgNP solution and shaken at 120 rpm. Every hour (for a total of 5 hours), one tube from each group was centrifuged at 4500 rpm to isolate the supernatant. Bacteria (and bound CC-AgNPs) were pulled down into the pellet while free CC-AgNPs (yellow in color) remained in the supernatant, which was measured using a spectrophotometer at 430 nm.

#### Simulation of WWTP Aeration Tanks

Three axles were connected using VEX Robotics chains and sprockets and moved with a VEX Robotics 393 Motor to ensure a constant RPM across all 3 axles (figure 5). Aeration tanks were simulated using clear plastic cylinders and the axles churned the mixture for 5 hours. To simulate the process in secondary sedimentation tanks, flocculant powder from Dihua WWTP (Taipei, Taiwan) was added to accelerate sedimentation after churning.

**Figure 5.**
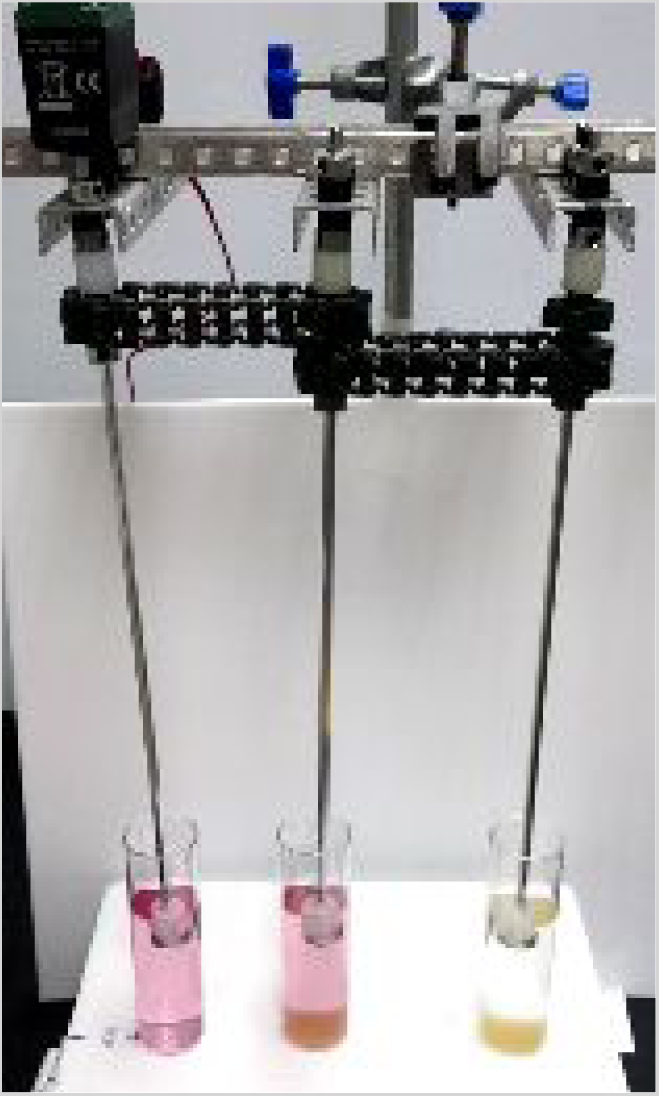
Setup of simulated WWTP tanks. Clear plastic cylinders and rotating central rods that extend into the cylinders mimic water movement in WWTP tanks.

### 2.2 Biofilm

#### Preliminary Biofilm and Gold Nanoparticle Assay

*E. coli* liquid cultures grown in LB were transferred to petri dishes, which contained glass coverslips to provide a surface for adherence. The dishes were incubated at 37°C for 7 to 14 days, with 2 mL of LB added every two days to prevent the media from drying out. Biofilms were extracted by pipette and washed with distilled water before use (figure 6C).

**Figure 6.**
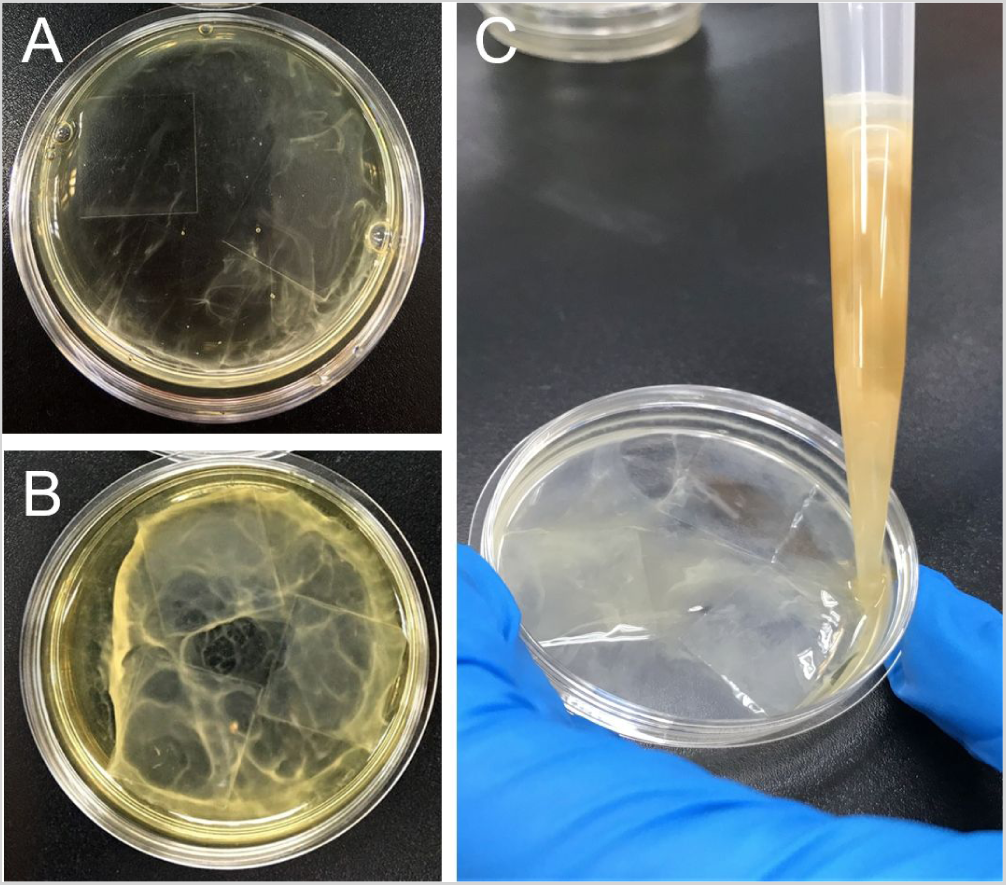
Growing *E. coli* biofilm. **A**,**B)** Liquid cultures were plated with glass coverslips and incubated for up to 2 weeks. **C)** Biofilm was washed with distilled water before use.

We tested biofilm’s ability to trap 30 nm gold nanoparticles (AuNPs; Sigma Aldrich, 741973). Four experimental groups were set up (figure 11A): a negative control containing only AuNPs, and three tubes containing AuNPs with either planktonic bacteria, biofilm, or “dead” biofilm (treated with ampicillin, chloramphenicol, and kanamycin). Samples were shaken for 24 hours and centrifuged to isolate the supernatant. Absorbance at 527 nm was measured to quantify the amount of free AuNPs (purple in color) in the supernatant. AuNP trapping would lead to a decrease in absorbance of the supernatant (figure 11B).

**Figure 7.**
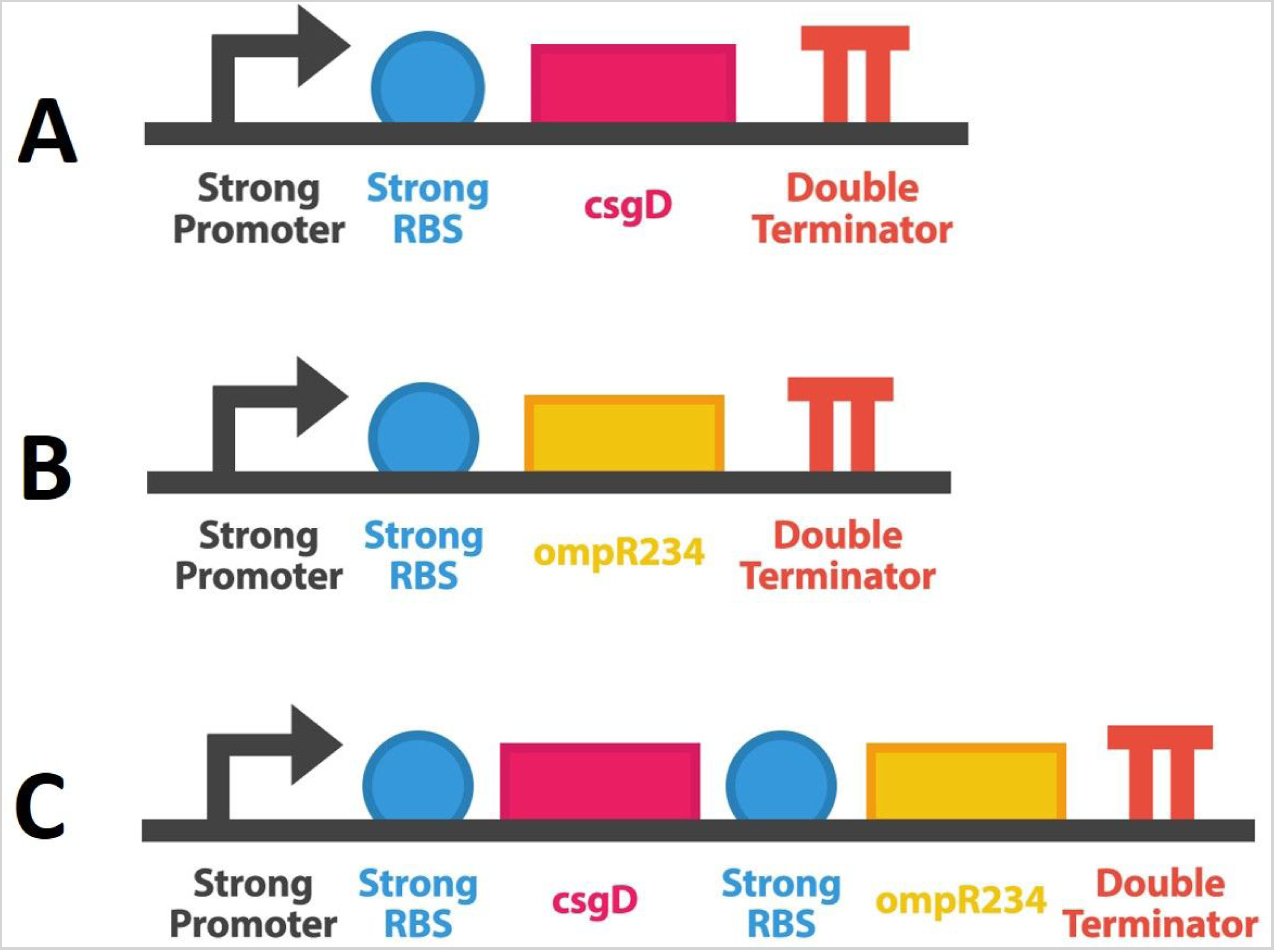
Construct designs to overexpress curli operon regulators. Overexpression of either CsgD (**A**, BBa_K2229100), OmpR234 (**B**, BBa_K2229200), or both proteins (**C**, BBa_K2229300).

**Figure 8.**
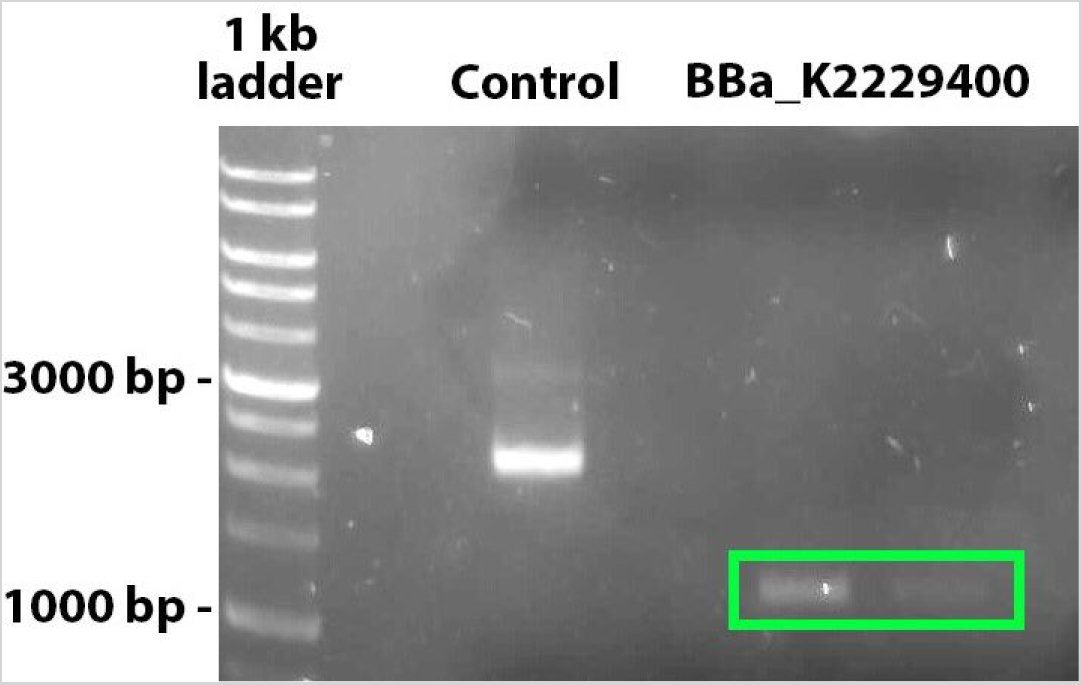
PCR check for the PR-expression construct (BBa_K2229400) using VF2 and VR primers. The expected size of BBa_K2229400 is 1300 bp (green box).

**Figure 9.**
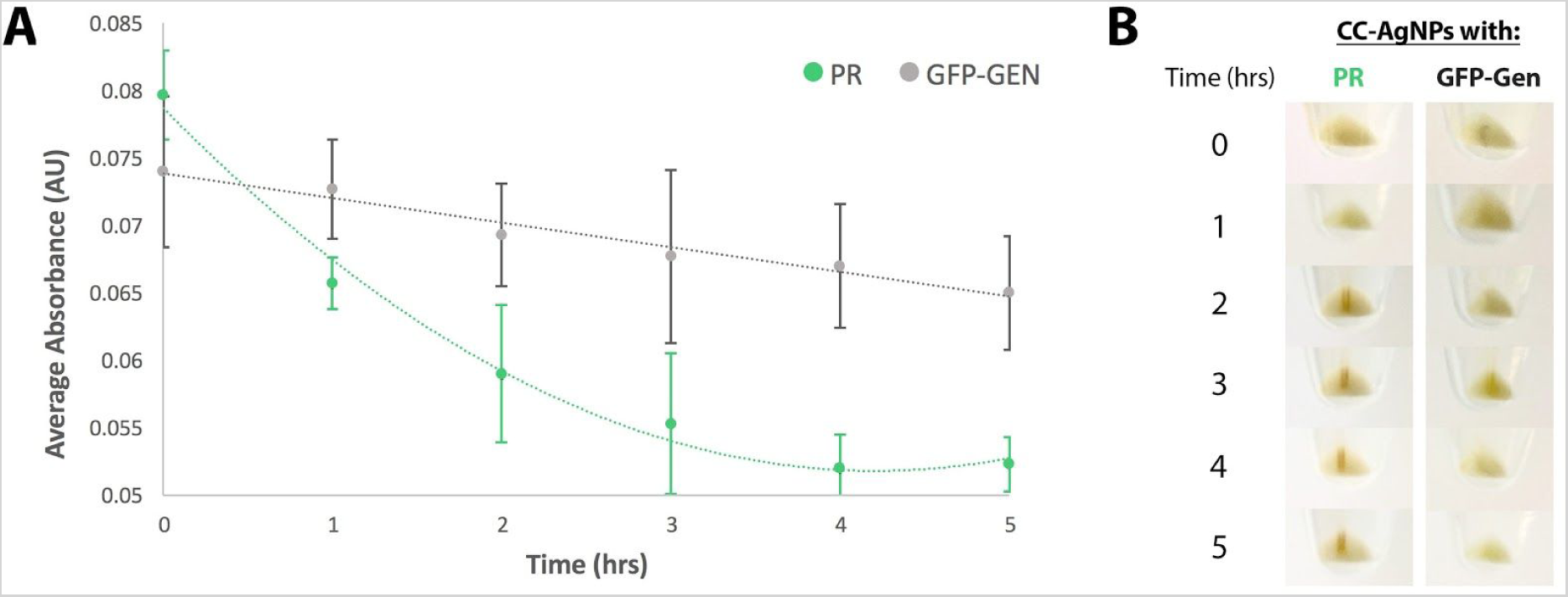
Proteorhodopsin traps CC-AgNPs. A) After 5 hours, absorbance of the supernatant significantly decreased when PR bacteria was added to CC-AgNPs, but not when GFP-Gen (negative control) bacteria was added. Error bars represent SEM. **B)** After centrifugation, we observed a large orange region (aggregated CC-AgNPs) in the PR bacteria pellet.

**Figure 10.**
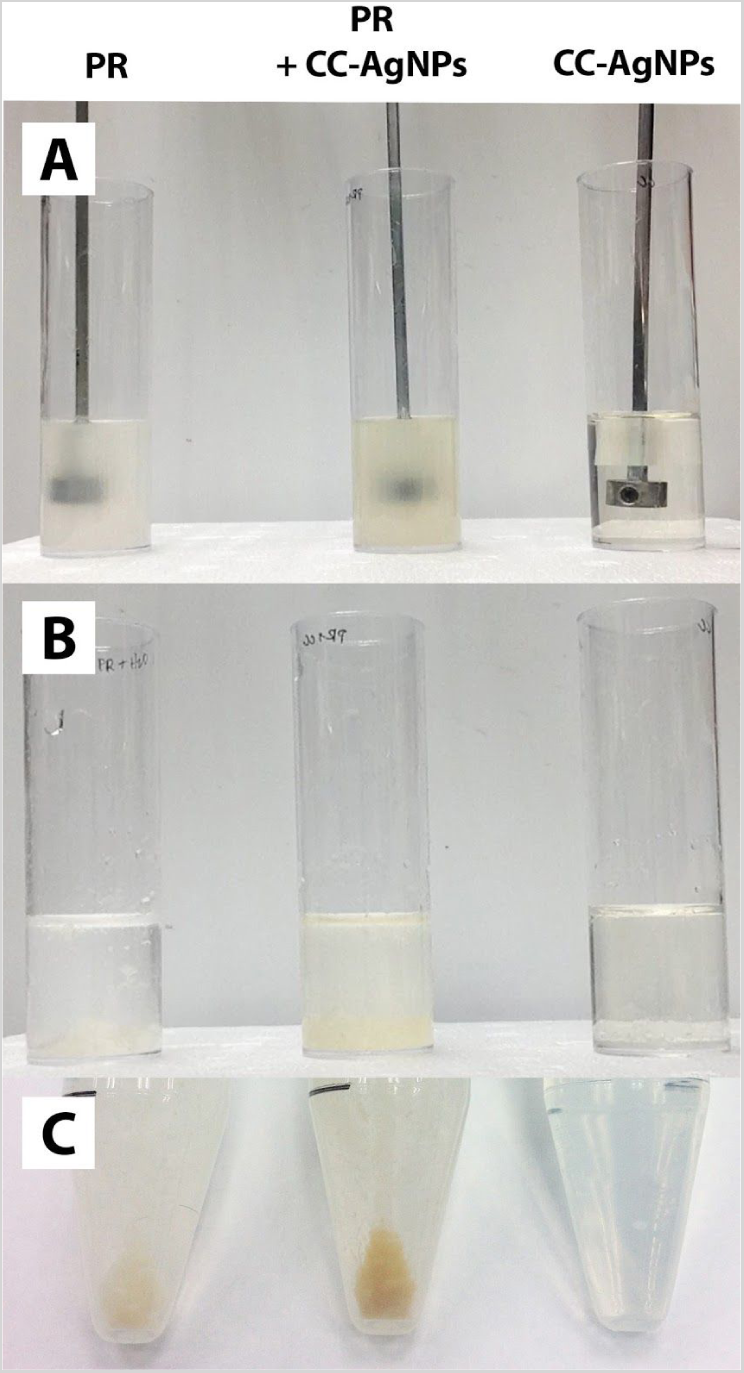
PR bacteria trap CC-NPs in simulated WWTP aeration tanks. A) Three groups were set up and churned for 5 hours: PR bacteria + distilled water, PR bacteria + CC-AgNPs, and CC-AgNPs + distilled water. **B)** After 5 hours, flocculants were added and aggregated materials settled to the bottom. **C)** The contents of each cylinder were centrifuged. The pellet of the PR bacteria + CC-AgNPs mixture (middle) was orange, reflecting the presence of aggregated CC-AgNPs.

**Figure 11.**
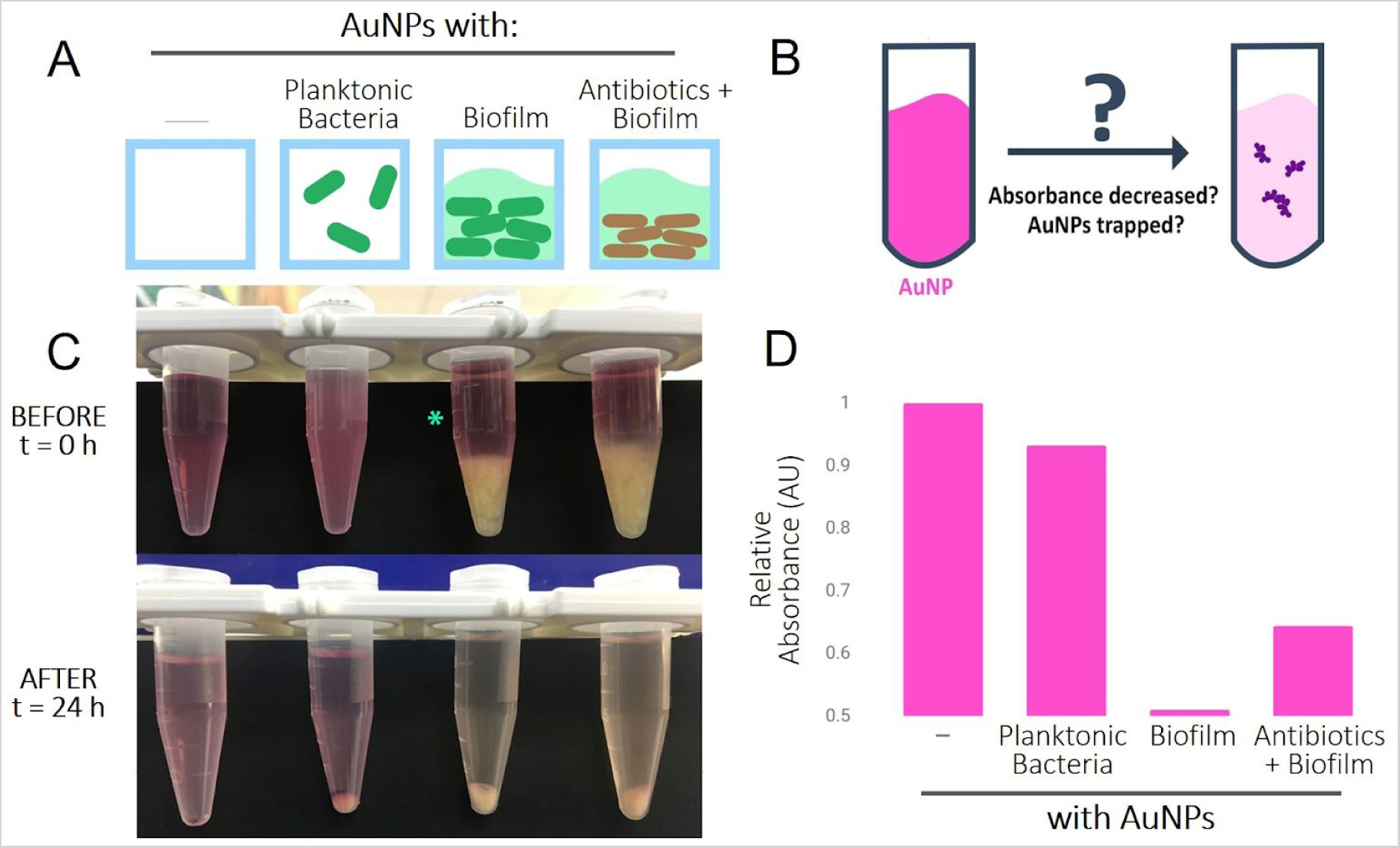
Biofilms trap NPs. A) Four experimental groups: AuNPs alone, or added to planktonic bacteria, biofilm, or antibiotic-treated biofilm. **B)** Absorbance of the solution (pink) should decrease if AuNPs become trapped. **C**,**D)** AuNPs were incubated with either bacteria or biofilm for 24 hours, and then centrifuged. Absorbance of each supernatant was measured, as free AuNPs should remain in the supernatant. Shown are representative images and graph.

#### Sample Preparation for SEM

Three different SEM sample fixation and preparation methods (table 1) were performed [27]. Image quality was further improved by taking multiple images of a field of view and stacking the images together to reduce noise.

**Table 1.** Comparison of three different SEM sample preparation protocols: critical point drying, fixation by glutaraldehyde (GA), and fixation by GA, Alcian blue, and potassium ferrocyanide.

| Method | Advantages | Disadvantages |
| --- | --- | --- |
| **Critical Point Drying (CPD)** | Fast | Deformed morphology<br>Loses EPS |
| **Fixation by GA** | Preserves cell structure | Loses EPS<br>Long fixation time |
| **Fixation by GA, Alcian blue, and $K_4Fe(CN)_6$** | Preserves cell & biofilm structure | Very long fixation time |

#### Construct Design, Assembly, and Expression

Three constructs were built to upregulate curli production by overexpressing CsgD, OmpR234 (a mutant form of OmpR which is constitutively active), or both (figure 7). We acquired all parts from the iGEM distribution kit: a strong promoter and strong RBS combination (BBa_K880005) to maximize protein production, strong RBS (BBa_B0034), *csgD* (BBa_K805015), *ompR234* (BBa_K342003), and a double terminator (BBa_B0015) to end transcription. Sequences were confirmed by Tri-I Biotech, and protein expression was confirmed by SDS-PAGE.

#### Congo Red Assay

Congo Red solution mixed with *E. coli* liquid cultures were transferred to 12-well microtiter plates, and incubated with glass coverslips at 37°C for one day. The samples were then washed with Phosphate Buffered Saline (PBS) and dried at 60°C. Stained biofilm on the glass coverslips appeared red, was solubilized in ethanol, and quantified by measuring the absorbance at 500 nm.

#### Simulation of WWTP Sedimentation Tank

Sedimentation tanks were similarly simulated using clear plastic cylinders (figure 5), with plastic biocarriers attached to a central spinning rotor. Biofilm was grown directly onto biocarriers in the cylinders to minimize any disturbances. A low rotation speed (60 rpm) was used to simulate the mild movement of water in sedimentation tanks.

## Results & Discussion

### 3.1 Proteorhodopsin

#### Proteorhodopsin Binds Citrate-Capped Nanoparticles

A PR-expressing construct, designed to target the citrate capping agent, was assembled (BBa_K2229400; figure 8). To test whether PR binds CC-NPs, we added CC-AgNPs to either *E.* c*oli* expressing PR or *E.* c*oli* not expressing PR (GFP-generator). We then centrifuged both groups and tracked the absorbance of the supernatant, which contains free NPs. Over 5 hours, we found that absorbance values of the supernatant at 430 nm decreased faster when PR bacteria was added, while the absorbance did not change significantly when GFP-generator bacteria was added (figure 9A). In addition, after centrifugation, we observed dark orange regions in the PR bacteria pellet, but not in the GFP-generator bacteria pellet (figure 9B). Since CC-AgNPs are yellow in color, the orange regions in the PR pellet are likely aggregated CC-AgNPs, showing that PR bacteria can bind CC-NPs to lower the levels of free CC-NPs.

#### Proteorhodopsin Bacteria Trap Citrate-Capped Nanoparticles in Simulated WWTP Tanks

We tested PR bacteria under conditions similar to a WWTP aeration tank. To simulate these conditions, we built our own “aeration tanks” using clear cylinders and a central rotor. Three groups were set up: PR bacteria alone, PR bacteria + CC-AgNP, or CC-AgNP solution alone (figure 10A). We turned on the rotor and churned the mixture for 5 hours. In the CC-AgNP cylinder, adding flocculants did not have any effect (figure 10B and C), suggesting that current wastewater treatment practices cannot remove NPs. In the cylinders containing PR bacteria, however, aggregated materials (including bacteria) settled to the bottom of the cylinder as expected (figure 10B). We then centrifuged the contents of each cylinder, and observed that the pellet of the PR bacteria + CC-AgNP mixture was orange, reflecting the presence of aggregated CC-AgNPs (figure 10C). In this WWTP aeration tank simulation, we show that PR bacteria bind to CC-AgNPs, and can be subsequently separated out by flocculation.

### 3.2 Biofilm

#### Biofilms Trap Nanoparticles

Using 30 nm AuNPs, we performed a preliminary test to measure biofilm’s ability to trap NPs (figure 11). Four experimental groups were set up: a negative control containing only AuNPs, and three tubes containing AuNPs with either planktonic bacteria, biofilm, or “dead” biofilm (treated with antibiotics) (figure 11A). If AuNPs were trapped, we would expect a decrease in absorbance (figure 11B). In the negative control and AuNP + planktonic bacteria group, the purple color of the AuNP solution did not change, indicating that bacteria alone cannot trap NPs (figure 11C,D). The addition of biofilm, however, greatly reduced the amount of AuNP in the supernatant (figure 11C,D). Even when bacteria in the biofilm were killed with antibiotics, AuNP levels in the supernatant were still reduced, suggesting that the removal of AuNPs depends on the sticky and slimy extracellular components of biofilm and not on the bacteria (figure 11C,D). When we fixed and imaged the biofilm + AuNP sample by SEM, we also observed NPs in EPS areas (figure 12). Together, our results suggest that the EPS layer of biofilms can trap NPs.

**Figure 12.**
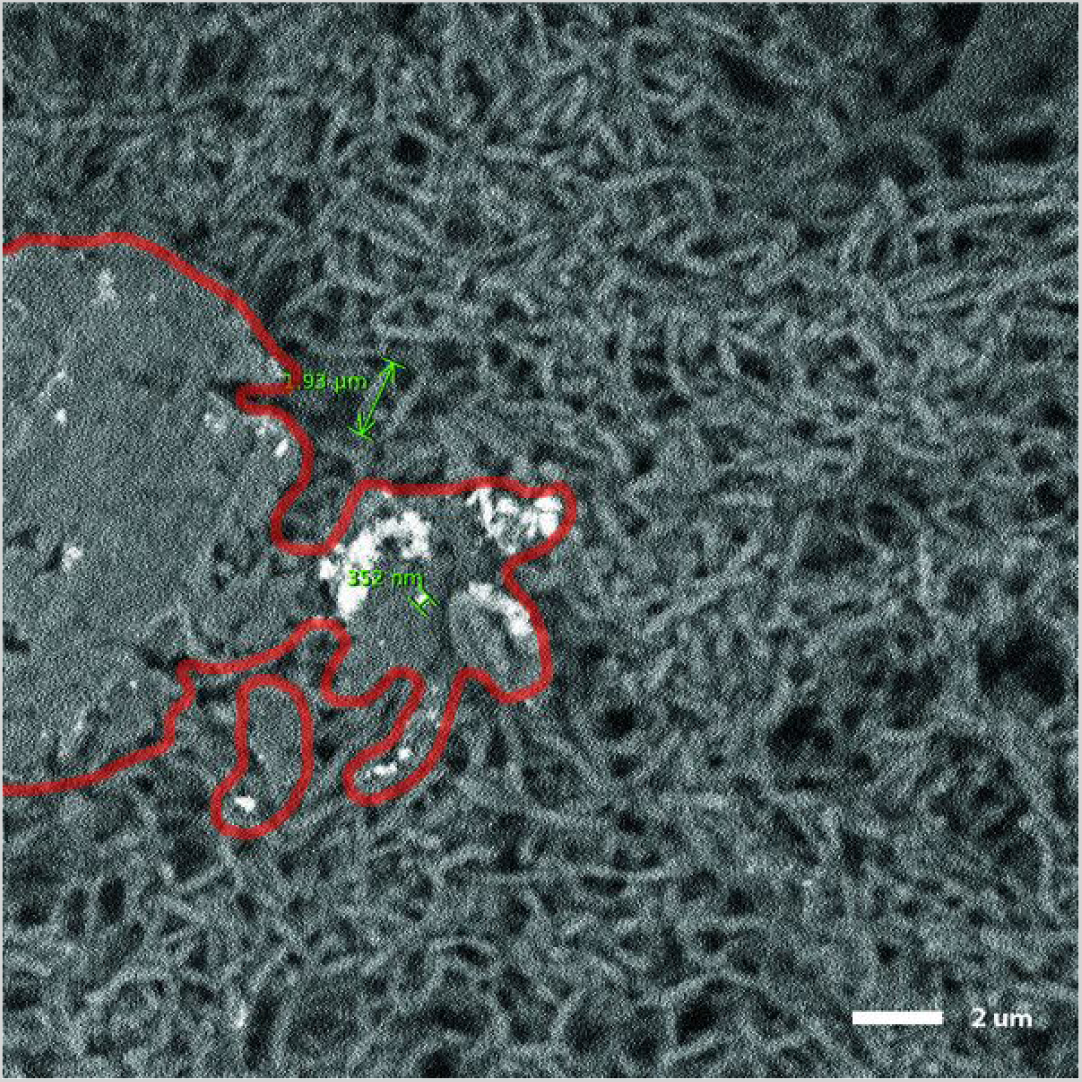
SEM image showing AuNPs trapped by biofilm. A biofilm + AuNP sample was fixed with GA, and imaged using SEM. Some EPS is preserved (outlined in red) and AuNPs (white) seemed to aggregate and adhere onto the EPS.

#### CsgD and OmpR234 Increases the Expression of Curli Operon Proteins

In *E. coli*, biofilm synthesis is mainly mediated by two curli operons [28]. Curli fibers are the main protein components of the EPS and promote biofilm formation by facilitating cell-surface and cell-cell adhesion [28, 29]. The operons (*csgBA* and *csgDEFG*) control the expression of six proteins essential to biofilm formation (figure 13). CsgA and CsgB are curli monomers which can polymerize to form curli fibers. CsgD is an activator of *csgBA* transcription, whereas CsgE, CsgF, and CsgG facilitate the extracellular transport of CsgA and CsgB. The operon *csgDEFG* can be activated by the protein OmpR, and the subsequent expression of all six proteins increases biofilm formation [28].

**Figure 13.**
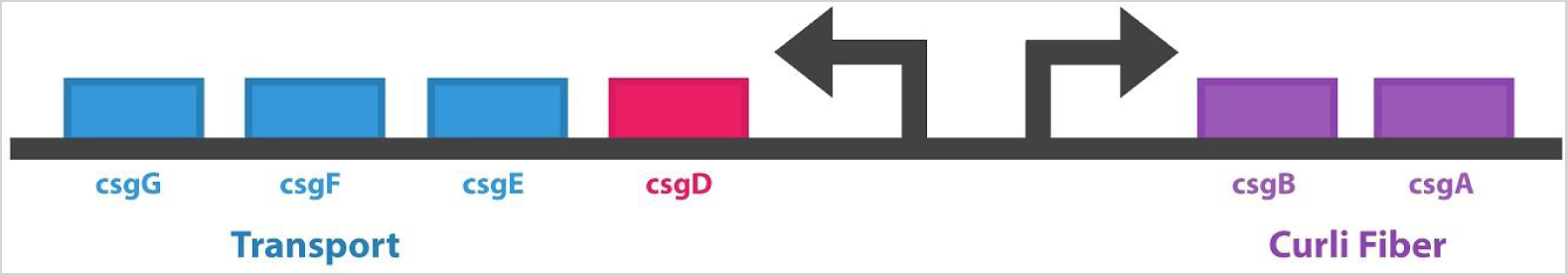
Two curli operons—csgBA and csgDEFG—direct biofilm synthesis. CsgD is an activator of csgBA transcription. The expressed CsgA and CsgB are curli monomers, which can be transported out of cells by the proteins CsgE, CsgF, and CsgG.

We hypothesized that biofilm production would be upregulated (in increasing order) if we overexpressed CsgD, OmpR234 (a constitutively active form of OmpR), or both (figure 14). Overexpression of CsgD should result in more curli monomers, but no transport proteins to carry those monomers out of the cell (figure 14A). Overexpression of OmpR234 should allow curli monomers to be exported and form curli fibers and biofilm (figure 14B). Finally, when both CsgD and OmpR234 are overexpressed, twice the amount of curli monomers should be made and exported to form even more curli fibers and biofilm (figure 14C).

**Figure 14.**
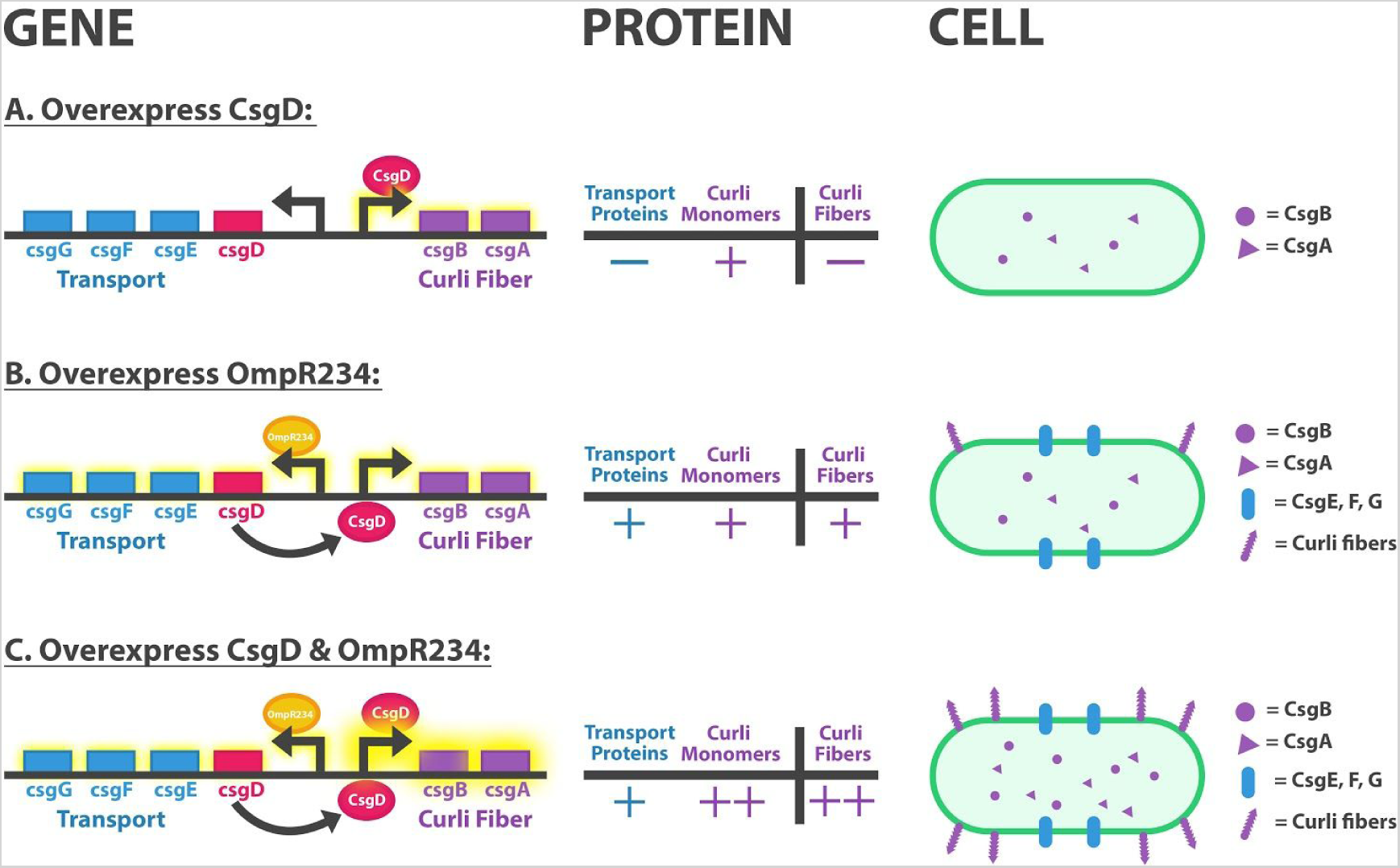
Overexpression of CsgD and/or OmpR234 should upregulate the curli operon to different degrees. We hypothesized that biofilm production would be upregulated (in increasing order) if we overexpressed **A)** CsgD, **B)** OmpR234, or **C)** both.

To achieve different levels of biofilm production, we designed and assembled constructs to only express CsgD (BBa_K2229100, figure 15), OmpR234 (BBa_K2229200, figure 15), or to express both regulator proteins (BBa_K2229300, figure 15). Next, we ran SDS-PAGE using transformed and lysed *E. coli* cultures to test the expression of CsgD and OmpR234 (figure 16). Cultures transformed with the basic parts BBa_K805015 (*csgD* ORF alone) and BBa_K342003 (*ompR234* ORF alone) were used as controls. We expected to see CsgD around 25 kDa and OmpR234 around 27 kDa [30, 31].

**Figure 15.**
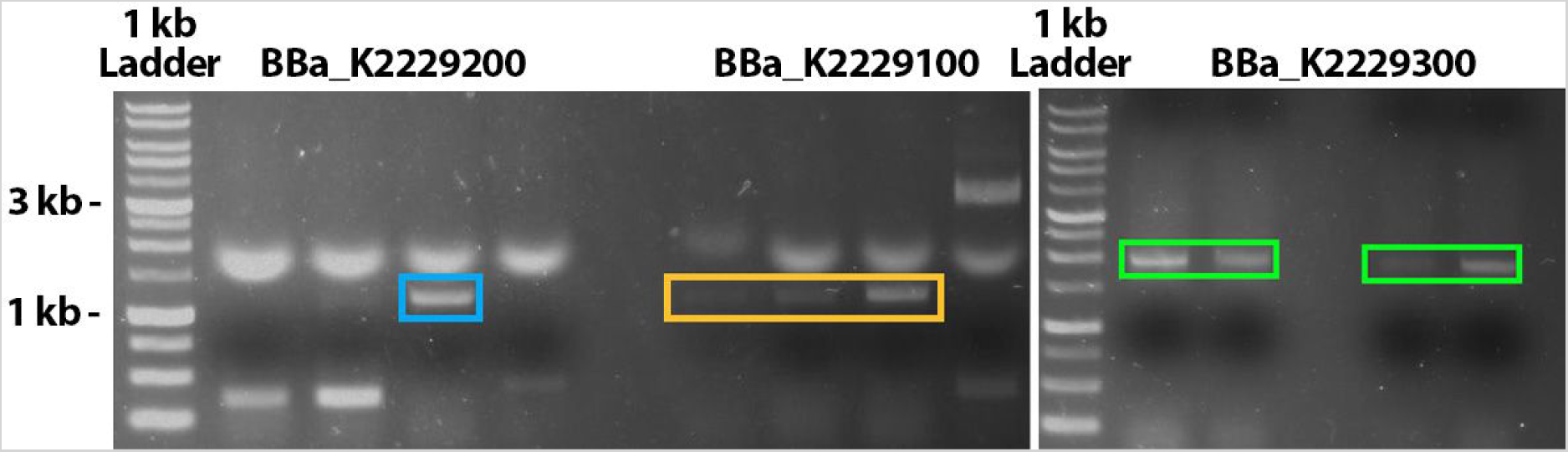
PCR check for the CsgD-expressing (BBa_K2229100), OmpR234-expressing (BBa_K2229200), and dual CsgD and OmpR234 expression (BBa_K2229300) constructs using VF2 and VR primers. The expected size of BBa_K2229100 is 1100 bp (orange box), BBa_K2229200 is 1200 bp (blue box), and BBa_K2229300 is 1900 bp (green boxes).

**Figure 16.**
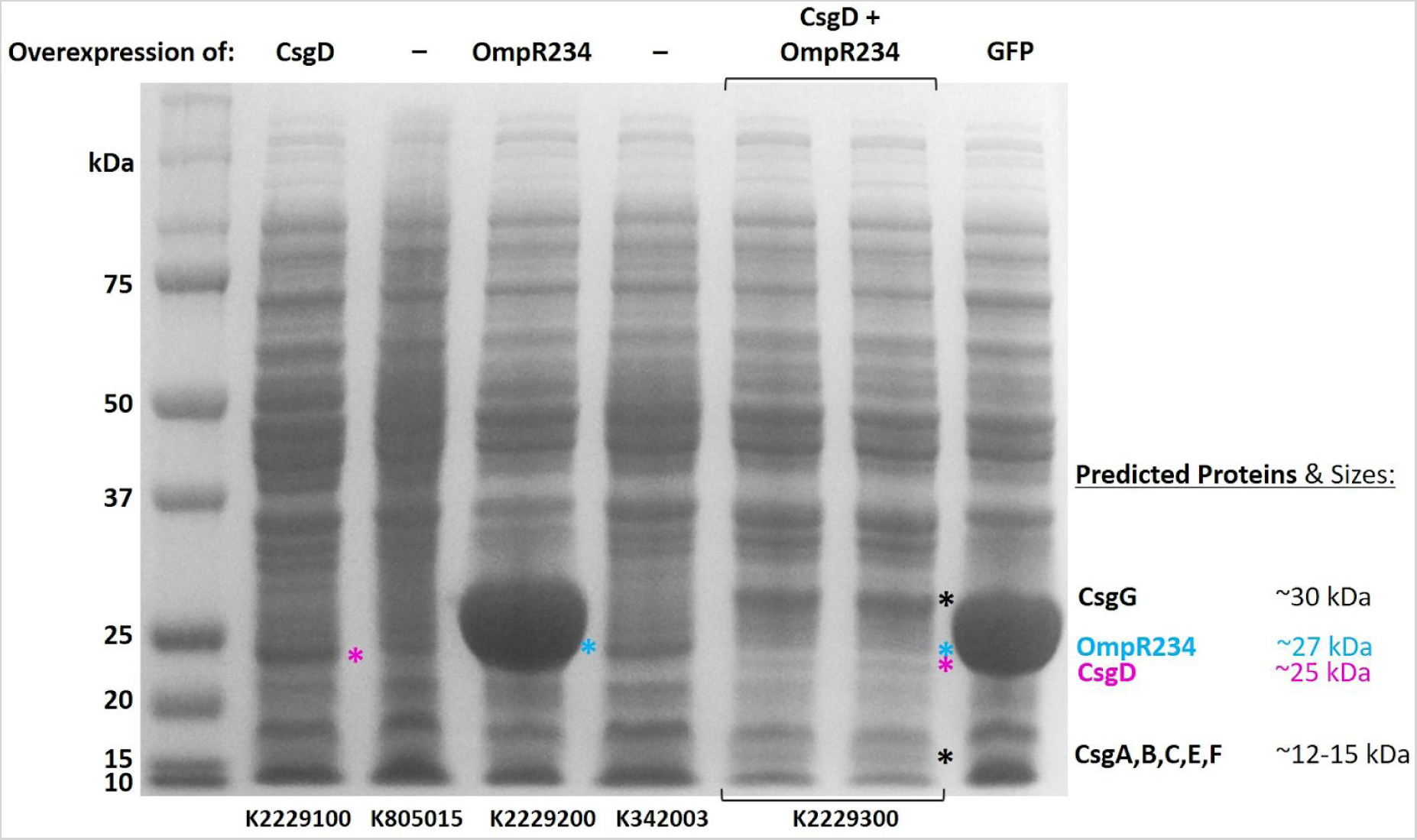
SDS-PAGE results show that BBa_K2229100, BBa_2229200, and BBa_K2229300 overexpress CsgD (pink asterisk), OmpR234 (blue asterisk), or both proteins, respectively. Other predicted expressed proteins from the curli operon (black asterisks) are listed on the right; *E. coli* expressing GFP was used as a control.

Compared to controls, thicker and darker bands at the expected sizes were observed for both BBa_K2229100 (CsgD overexpression) and BBa_K2229200 (OmpR234 overexpression) (figure 16; proteins bands are marked by asterisks). In addition to the bands at 25 and 27 kDa, cultures carrying BBa_K2229300 (both CsgD and OmpR234 expression) contained two extra bands at 15 kDa and 30 kDa, which were not observed in the controls. We looked into the other curli proteins and found that CsgG is around 30 kDa, whereas CsgA, B, C, E, and F are all around 15 kDa [32, 33, 34]. This suggests that, as hypothesized, BBa_K2229300 stimulates the expression of other curli proteins (predicted proteins and sizes are labeled in figure 16).

#### CsgD and OmpR234 Increases Biofilm Production

Seeing that overexpression of CsgD and OmpR234 stimulates other curli proteins as well (figure 16), we next tested if our constructs actually yield greater biofilm production. Bacteria carrying different constructs were cultured using 12-well microtiter plates, and stained with Congo Red, a dye commonly used to measure biofilm production [35]. Wells with higher biofilm production would retain more stain, which we could quantify by taking absorbance measurements.

Our results show that overexpressing CsgD and/or OmpR234 increased biofilm production to different degrees, as hypothesized (figure 14). Overexpression of both OmpR234 and CsgD (BBa_K2229300) increased biofilm production the most (figure 17). BBa_K2229300 also increased adhesion to glass coverslips placed inside the wells, and we observed a layer of biofilm which remained attached to the glass surface after the washing steps (figure 17A).

**Figure 17.**
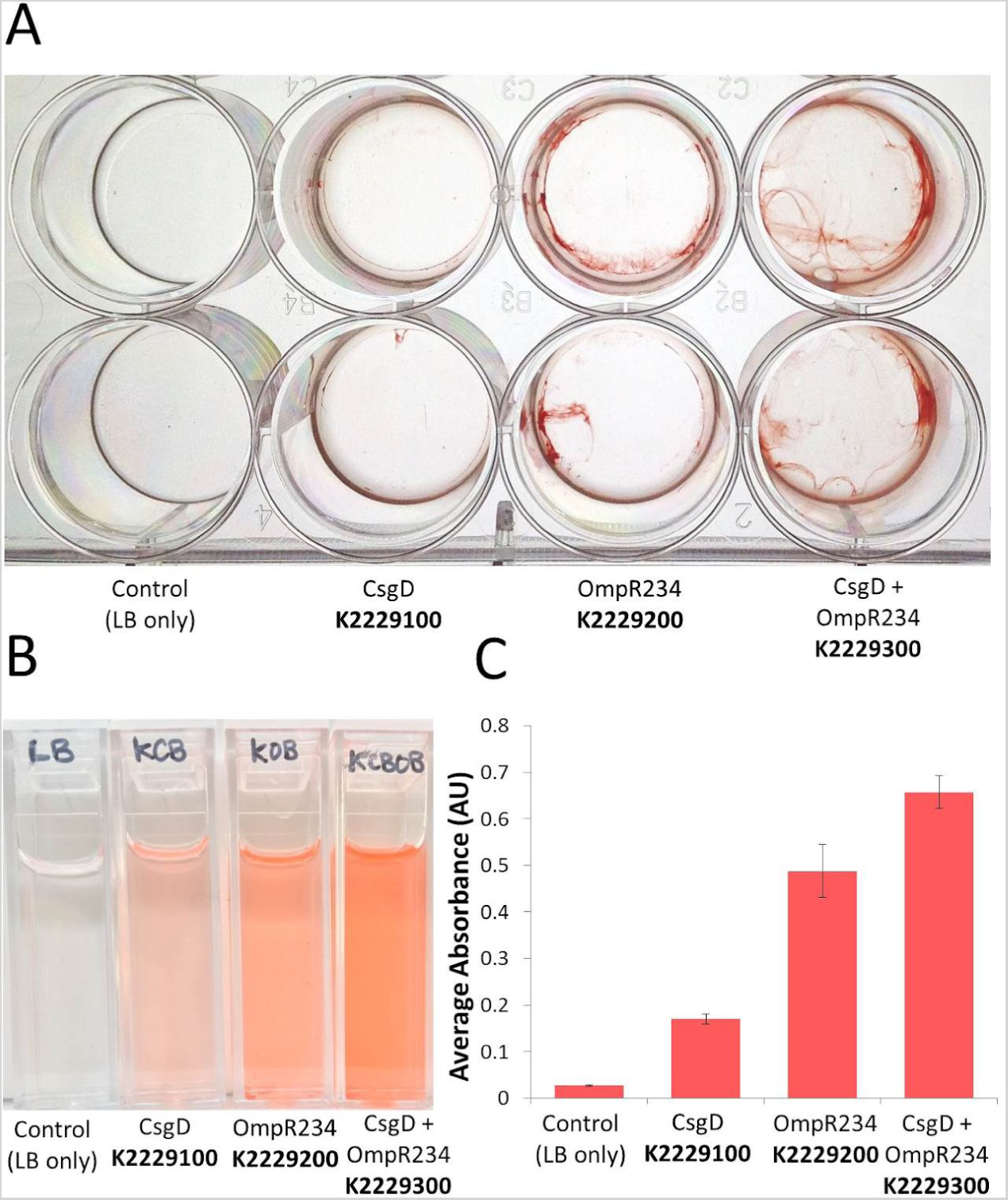
Overexpression of CsgD and OmpR234 increases biofilm production to different degrees. **A)** Congo Red dye stains biofilms. Biofilm production increased as CsgD, OmpR234, or both proteins combined were overexpressed. BBa_K2229300 also increased adhesion to the glass coverslip surface. **B)** Stained biofilm was solubilized in ethanol, and absorbance was measured at 500 nm **C)**. Error bars represent SEM.

#### Biofilm Traps AuNPs in Simulated WWTP Tanks

To simulate the implementation of biofilms in a WWTP secondary sedimentation tank, we used clear plastic cylinders with biocarriers attached to a central spinning rotor. Three groups were set up: biofilm alone, biofilm + AuNP, and AuNP solution alone (figure 18). Biofilms were added directly into the cylinders, so they could attach to the biocarriers in the simulated tanks.

**Figure 18.**
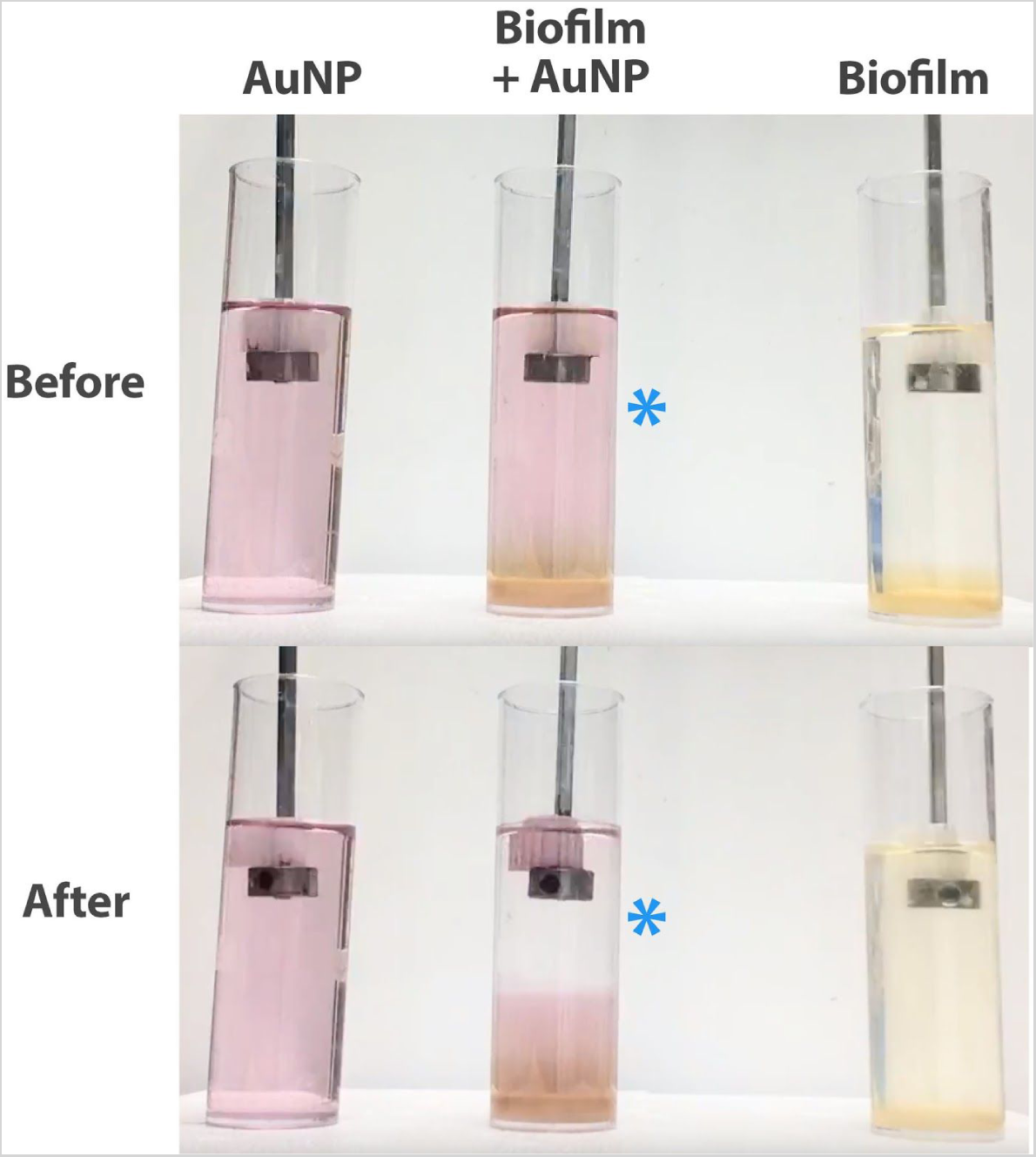
Biofilms effectively remove NPs in a simulated sedimentation tank. After about 30 hours of mixing, the purple color of the AuNP solution turned clear (blue asterisk) in the biofilm + AuNP group (middle). AuNP alone (left) and biofilm alone (right) were used as controls.

After about 30 hours of mixing, the purple color of the biofilm + AuNP solution turned clear, and we observed purple aggregates forming on the rotating biocarriers (figure 18). In contrast, the AuNP alone group did not change at all (figure 18 left). These results suggest that biofilms were able to first attach to biocarriers, then effectively remove NPs in a simulated secondary sedimentation tank.

## Conclusion

The recent rise in commercial NP usage and its potential health and environmental risks call for effective NP cleanup in WWTPs. Here, we show that the membrane protein PR binds to citrate, the most commonly used NP capping agent. This could be implemented in WWTP aeration tanks, where microbes are already present in the treatment process. In addition, we show that overexpressing biofilm regulators increases *E. coli* biofilm production, which can remove NPs. This biofilm can be coated onto biocarriers and used in WWTP sedimentation tanks. We envision using these two approaches to maximize NP removal in WWTPs.

## Acknowledgments

We would like to thank Mr. Matthew Fagen for providing assistance and materials for our biocarrier designs and WWTP model, Dr. Nicholas Ward for input on nanoparticle interaction with our constructs, Dr. Gwo-Dong Roam for sharing his insight on nanoparticle pollution, and Dihua WWTP (Taipei, Taiwan) and Boswell WWTP (Pennsylvania, USA) for answering our questions regarding current wastewater treatment methods.

## References

1. Vert M, Doi Y, Hellwich KH, Hess M, Hodge P, Kubisa P, et al. Terminology for biorelated polymers and applications (IUPAC Recommendations 2012). Pure Appl Chem. 2012; 84(2): 377–410. doi:10.1351/PAC-REC-10-12-04.

2. Vance ME, Kuiken T, Vejerano EP, McGinnis SP, Hochella MF, Rejeski D, et al. Nanotechnology in the real world: Redeveloping the nanomaterials consumer products inventory. Beilstein J Nanotechnol. 2015; 6: 1769–80. doi:10.3762/bjnano.6.181.

3. Ahamed M, Alsalhi MS, Siddiqui M. Silver nanoparticle applications and human health. Clinica Chimica Acta. 2010; 411(23-24): 1841–8. doi:10.1016/j.cca.2010.08.016

4. Kulthong K, Srisung S, Boonpavanitchakul K, Kangwansupamonkon W, Maniratanachote R. Determination of silver nanoparticle release from antibacterial fabrics into artificial sweat. Part Fibre Toxicol. 2010; 7: 8. doi:10.1186/1743-8977-7-8.

5. Lewicka ZA, Yu WW, Oliva BL, Contreras EQ, & Colvin VL. Photochemical behavior of nanoscale TiO2 and ZnO sunscreen ingredients. J. Photochem. Photobiol. 2013; 263, 24–33. doi:10.1016/j.jphotochem.2013.04.019

6. Ivask A, Kurvet I, Kasemets K, Blinova I, Aruoja V, Suppi S, et al. Size-dependent toxicity of silver nanoparticles to bacteria, yeast, algae, crustaceans and mammalian cells in vitro. PLoS One. 2014; 9(7):e102108. doi:10.1371/journal.pone.0102108.

7. Bondarenko O, Juganson K, Ivask A, Kasemets K, Mortimer M, Kahru A. Toxicity of Ag, CuO and ZnO nanoparticles to selected environmentally relevant test organisms and mammalian cells in vitro: a critical review. Archives of Toxicology. 2013; 87(7):1181–1200. doi:10.1007/s00204-013-1079-4.

8. Yin L, Colman BP, McGill BM, Wright JP, Bernhardt ES. Effects of silver nanoparticle exposure on germination and early growth of eleven wetland plants. PLoS One. 2012; 7(10):e47674. doi:10.1371/journal.pone.0047674.

9. Laban G, Nies L, Turco R, Bickham J, Sepulveda M. The effects of silver nanoparticles on fathead minnow (Pimephales promelas) embryos. Ecotoxicology. 2009; 19. 185-95. 10.1007/s10646-009-0404-4.

10. Paddle-Ledinek JE, Nasa Z. Effect of Different Wound Dressings on Cell Viability and Proliferation. Plast. Reconstr. Surg. 2006; 117(7S), 110–118. doi:DOI: 10.1097/01.prs.0000225439.39352.ce

11. Xu F, Piett C, Farkas S, Qazzaz M, Syed NI. Silver nanoparticles (AgNPs) cause degeneration of cytoskeleton and disrupt synaptic machinery of cultured cortical neurons. Mol. Brain Res. 2013; 6, 29. http://doi.org/10.1186/1756-6606-6-29

12. Mueller NC, Nowack B. Exposure modeling of engineered nanoparticles in the environment. Environ Sci Technol. 2008; 42(12):4447–53. doi:10.1021/es7029637.

13. Holder AL, Vejerano EP, Zhou X, Marr LC. Nanomaterial disposal by incineration. Environ Sci Process Impacts. 2013; 15(9):1652–64. doi:10.1039/c3em00224a.

14. Pescod, MB. Wastewater treatment and use in agriculture Rome: United Nations. 1992; 47

15. Davis PS. The Biological Basis of Wastewater Treatment. Strathkelvin Instruments. 2005. Available from:www.s-can.nl/media/1000154/thebiologicalbasisofwastewatertreatment.pdf.

16. Kiser MA, Westerhoff P, Benn T, Wang Y, Pérez-Rivera J, Hristovski K. Titanium nanomaterial removal and release from wastewater treatment plants. Environ Sci Technol. 2009; 43(17):6757–63. doi:10.1021/es901102n.

17. Levard C, Hotze EM, Lowry GV, Brown GE. Environmental transformations of silver nanoparticles: impact on stability and toxicity. Environ Sci Technol. 2012; 46(13):6900–14. doi:10.1021/es2037405.

18. Syed FF. Citrate Binding to the Membrane Protein Proteorhodopsin. Ph.D. Dissertation, Syracuse University. 2011. Available from: https://surface.syr.edu/che_etd/182/

19. Sehar S, Naz I. Role of the Biofilms in Wastewater Treatment. In: Dhanasekaran D, Thajuddin N, editors. Microbial Biofilms - Importance and Applications. InTech; 2016: pp.121–143. doi:10.5772/63499.

20. Yun J, Lee DG. Silver Nanoparticles: A Novel Antimicrobial Agent. In: Grumezescu AM, editors. Antimicrobial Nanoarchitectonics; 2017: pp.139–166. doi:10.1016/b978-0-323-52733-0.00006-9

21. Walden C, Zhang W. Biofilms Versus Activated Sludge: Considerations in Metal and Metal Oxide Nanoparticle Removal from Wastewater. Environ Sci Technol. 2016; 50(16):8417–31. doi:10.1021/acs.est.6b01282.

22. Donlan RM. Biofilms: Microbial life on surfaces. Emerg Infect Dis. 2002; 8(9):881–890. doi:10.3201/eid0809.020063

23. Nevius BA, Chen YP, Ferry JL, Decho AW. Surface-functionalization effects on uptake of fluorescent polystyrene nanoparticles by model biofilms. Ecotoxicology. 2012; 21(8):2205–13. doi:10.1007/s10646-012-0975-3.

24. Choi O, Yu C, Esteban Fernández G, Hu Z. Interactions of nanosilver with Escherichia coli cells in planktonic and biofilm cultures. Water Res. 2010; 44(20):6095–6103. doi:10.3201/eid0809.020063.

25. Fattahi S, Kafil HS, Nahai MR, Asgharzadeh M, Nori R, Aghazadeh M. (2015). Relationship of biofilm formation and different virulence genes in uropathogenic Escherichia coli isolates from Northwest Iran. GMS Hyg Infect Control. 2015; 10: Doc11. doi:10.3205/dgkh000254.

26. United States Environmental Protection Agency. Final risk assessment of Escherichia coli K-12 derivatives. EPA. 1997. Available from: https://www.epa.gov/sites/production/files/2015-09/documents/fra004.pdf.

27. Fischer, ER, Hansen, BT, Nair V, Hoyt FH, Dorward DW. Scanning electron microscopy. Curr Protoc Microbiol. 2012; Chapter 2: Unit 2B.2. doi:10.1002/9780471729259.mc02b02s25.

28. Barnhart M, Chapman M. Curli Biogenesis and Function. Annual Review of Microbiology. 2006; 60(1):131–147. doi:0.1146/annurev.micro.60.080805.142106.

29. Reichhardt C, Jacobson AN, Maher MC, Uang J, McCrate OA, Eckart M, et al. Congo Red Interactions with Curli-Producing E. coli and Native Curli Amyloid Fibers. PLoS One. 2015; 10(10):e0140388. doi:10.1371/journal.pone.0140388.

30. Brombacher E, Baratto A, Dorel C, Landini P. Gene expression regulation by the Curli activator CsgD protein: modulation of cellulose biosynthesis and control of negative determinants for microbial adhesion. J Bacteriol. 2006; 188(6):2027–2037. doi:10.1128/JB.188.6.2027-2037.2006.

31. Martinez-Hackert E, Stock AM (1997). The DNA-binding domain of OmpR: crystal structures of a winged helix transcription factor. Structure. 1997; 5(1):109–24. doi:10.1016/S1074-5521(97)90269-6.

32. Robinson LS, Ashman EM, Hultgren SJ, Chapman MR. Secretion of curli fibre subunits is mediated by the outer membrane-localized CsgG protein. Mol Microbiol. 2006; 59(3):870–81. doi:10.1111/j.1365-2958.2005.04997.x.

33. Uhlich GA, Gunther NW, Bayles DO, Mosier DA. The CsgA and Lpp proteins of an Escherichia coli O157:H7 strain affect HEp-2 cell invasion, motility, and biofilm formation. Infect Immun. 2009; 77(4):1543–52. doi:10.1128/IAI.00949-08.

34. Shu Q, Crick SL, Pinkner JS, Ford B, Hultgren SJ, Frieden C. The E. coli CsgB nucleator of curli assembles to β-sheet oligomers that alter the CsgA fibrillization mechanism. Proc Natl Acad Sci USA. 2012; 109(17):6502–7. doi:10.1073/pnas.1204161109.

35. Reinke AA, Gestwicki JE. Insight into Amyloid Structure Using Chemical Probes. Chem Bio Drug Des. 2011; 77(6), 399–411.http://doi.org/10.1111/j.1747-0285.2011.01110.x

