## Supplementary Materials for "Recombinant expression of Proteorhodopsin and biofilm regulators in *Escherichia coli* for nanoparticle binding and removal in a wastewater treatment model"

- Nathaniel Gore

  Hi TAS-Taipei! Thanks for your submission - we moved it into the PLOS Submission room so people would be able to discover it. I've added Justin Wang as a co-editor.

  3 days ago

  Reply Like

- Nathaniel Gore

  If there are any supplementary information files - data tables for example - please can you make them available to readers.

  3 days ago

  Reply Like

- Nathaniel Gore

  Our editors suggest that it would be useful to compare your system to the others already published, with a clear discussion of limitations, advantages, sensitivity and costs.

  3 days ago

  Reply Like

- Nathaniel Gore

  We also suggest providing details of the cell line sources and culture conditions to ensure your work is reproducible.

  3 days ago

  Reply Like

- Zi Fei Wang

  Hello! I would love to learn more about the Human Practices aspects of your project! This includes: was there a reason your team chose synthetic biology as the solution for the production of proteorhodopsin through E. Coli? Are there current methods for nanoparticle removal in wastewater treatment? If yes, it may be beneficial to compare your synbio method to the procedures currently employed.
