## Supplementary Materials for "Recombinant expression of Proteorhodopsin and biofilm regulators in *Escherichia coli* for nanoparticle binding and removal in a wastewater treatment model"

**Comments 1 and 3:**
**Compare your system to the others already published, with a clear discussion of limitations, advantages, sensitivity and costs?**

**Are there any current methods for nanoparticle removal in wastewater treatment? If yes, it may be beneficial to compare your synbio method to the procedures currently employed.**

A vast majority of municipal wastewater treatment plants (WWTPs) do not have specialized procedures designed to remove NPs in wastewater. Instead, NPs are subject to the conventional procedures used to treat larger particulates. When wastewater enters a WWTP, the first step is to remove coarse solids and large materials using a grit screen (figure 5-1). The water can then be processed in three main stages: Primary, Secondary, and sometimes Tertiary Treatment (Pescod 1992). In Primary Treatment, heavy solids are removed by sedimentation while floating materials (such as oils) can be taken out by skimming. However, dissolved materials and colloids—small, evenly dispersed solids such as NPs—are not removed here (Pescod 1992). Secondary Treatment generally involves the use of aeration tanks, where aerobic microbes help to break down organic materials. This is also known as the activated sludge process (Davis 2005). While this process removes some NPs, complete NP removal has not yet been achieved. A study monitoring an Arizona WWTP found that while sedimentation is effective at filtering large aggregates (72% removal rate), most small-sized $TiO_2$ NPs (41% removal rate) can still pass through the WWTP and enter major water systems downstream (Kiser et al. 2009). In certain WWTPs, wastewater may go through Tertiary Treatment, an advanced process typically aimed to remove nitrogen and phosphorous, and assumed to produce an effluent free of viruses (Pescod 1992). However, Tertiary Treatment requires additional infrastructure that is expensive and complex, limiting its global usage (Pescod 1992; Malik 2014).

One major advantage of our proposed system to remove NPs is that it can be used in any conventional WWTP with a secondary treatment step, where microbes are already used to break down organic matter. Our approach would involve adding two types of bacteria to an existing system, so the additional operational costs should be minimal.

There are also other advantages of using biological treatment over chemical or mechanical methods (like flocculant/coagulant polymers, filtration, or reverse osmosis). Chemical or mechanical processes generally require expensive infrastructure, is energy intensive, or is non-renewable (Pescod 1992; Malik 2014). Biological treatment, on the other hand, is more energy-efficient (the microbes will naturally break down or capture waste particles) and renewable. For example, once added to aeration tanks, the Proteorhodopsin (PR)-producing *E. coli* should be able to multiply and maintain its population. The same is true for biofilm-producing *E. coli*, which would be added to sedimentation tanks.

The use of two different types of bacteria also increases the chances of NP removal. Our PR *E. coli,* added to the aeration tanks, specifically targets citrate, the most common coating for NPs used in consumer products. Before wastewater enters the following sedimentation step, our PR *E. coli* should have already removed a portion of NP waste. Remaining NPs can then be captured by biofilm in the following sedimentation tank.

We tested the efficacy of the PR *E. coli* and biofilm at removing different types of NPs under simulated tank conditions. In our model, we found that our PR bacteria captured around 34% of citrate-capped AgNPs after a typical tank retention time of 5 hours. For the biofilm approach, we also show that our biofilm can attach onto a surface and start trapping 30nm AuNPs. In our simulation, we saw near-complete removal when we used 30mL of a 34uM AuNP solution--we observed the solution turn from purple (color of AuNP) to clear. Even with these simulations, however, we have not tested our bacteria in real WWTPs under actual sewage conditions, so it's hard to say how effective our system will be in real life.

**References:**

Davis, Peter S. "The Biological Basis of Wastewater Treatment." s-Can.nl, 2005, www.s-can.nl/media/1000154/thebiologicalbasisofwastewatertreatment.pdf.

M. A. Kiser, P. Westerhoff, T. Benn, Y. Wang, J. Pérez-Rivera, and K. Hristovski. (2009). Titanium Nanomaterial Removal and Release from Wastewater Treatment Plants. Environmental Science & Technology, 43 (17), 6757-6763

Malik, O. (2014, January 22). Primary vs. Secondary: Types of Wastewater Treatment. Retrieved October 12, 2017, from http://archive.epi.yale.edu/case-study/primary-vs-secondary-types-wastewater-treatment

Pescod, M. (1992). Wastewater treatment and use in agriculture (Vol. 47). Rome: United Nations. Vert, M., Doi, Y., Hellwich, K., et al. (2012). Terminology for biorelated polymers and applications (IUPAC Recommendations 2012). Pure and Applied Chemistry, 84(2), pp. 377-410. Retrieved 9 Oct. 2017, from doi:10.1351/PAC-REC-10-12-04

**Comment 2:**
**We also suggest providing details of the cell line sources and culture conditions to ensure your work is reproducible.**

**Cell line sources:**
We worked with *E. coli* K-12 (DH5-alpha).
All plasmids and DNA parts were obtained from the iGEM Parts Registry.

**PR Culture Conditions + OD Standardization**
PR-expressing *E. coli* liquid cultures grown in 3mL LB broth (Sigma Aldrich) and 25µg/mL chloramphenicol. Liquid cultures were shaken at 37˚C (180 rpm) overnight.

To isolate and standardize bacteria populations, each liquid culture was centrifuged at 4500 rpm for 2 minutes, and the pellet was resuspended in 3mL distilled water. This was repeated one more time and resuspended in 6mL distilled water. We measured OD600 for each sample using a UV-vis spectrophotometer (calibrated with distilled water). To standardize bacteria populations, all samples were diluted to match the lowest bacteria population.

### Biofilm Culture Conditions for Congo Red Assays

Liquid cultures of *E. coli* with our biofilm constructs were grown in 3mL LB (Sigma Aldrich) and 25µg/mL chloramphenicol. Liquid cultures were shaken overnight at 37˚C (180 rpm). 1mL culture was then transferred into a single well of a 12-well microtiter plate, containing a circular glass coverslip (20mm diameter) to provide a surface for adherence. For the Congo Red assay, 0.01g Congo Red was added to the same well. The plate was then incubated for 1 day at 37˚C in a stationary incubator.

**Comment 3:**
**Was there a reason your team chose synthetic biology as the solution for the production of proteorhodopsin through E. coli?**

Synthetic biology allows us to easily control the expression levels (for instance, by changing the promoters and ribosome binding sites) and modify the sequence of proteorhodopsin. Using synthetic biology, we could also express Proteorhodopsin protein in *E. coli,* one of the most common aerobic microbes found in wastewater aeration tanks. This allowed us to integrate our specific approach for targeting citrate-capped NPs into existing WWTPs.
